# A Deep Learning Framework for Predicting Prognostically Relevant Consensus Molecular Subtypes in HPV-Positive Cervical Squamous Cell Carcinoma from Routine Histology Images

**DOI:** 10.1101/2024.08.16.608264

**Authors:** Ruoyu Wang, Gozde N. Gunesli, Vilde Eide Skingen, Kari-Anne Frikstad Valen, Heidi Lyng, Lawrence S. Young, Nasir Rajpoot

## Abstract

Despite efforts in human papillomavirus (HPV) prevention and screening, cervical cancer remains the fourth most prevalent cancer among women globally. In this study, we propose an end-to-end deep learning framework to investigate histological correlates of the two consensus molecular subtype (CMS) of HPV-positive cervical squamous cell carcinoma (CSCC) patients. Analysing three international CSCC cohorts (n=545 patients), we demonstrate that the genomically determined CMS can be predicted from routine haematoxylin and eosin (H&E)-stained histology slides, with our Digital-CMS scores achieving significant patient stratifications in terms of disease-specific survival (TCGA p=0.0022, Oslo p=0.0495) and disease-free survival (TCGA p=0.0495, Oslo p=0.0282). In addition, our extensive analyses reveal distinct tumour microenvironment (TME) differences between the two CMS subtypes of the CSCC cohorts. Notably, CMS-C1 CSCC subgroup has markedly increased lymphocyte presence, whereas CMS-C2 subgroup has high nuclear pleomorphism, an elevated neutrophil-to-lymphocyte ratio, and increased neutrophil density. Analysis of representative histological regions reveals higher degree of malignancy in CMS-C2 patients, associated with poor prognosis. This study introduces a potentially clinically advantageous Digital-CMS score derived from digitised WSIs of routine H&E-stained tissue sections, offers new insights into TME differences impacting patient prognosis and potential therapeutic targets, and identifies histological patterns serving as potential surrogate markers of the two CMS subtypes for clinical application.

## INTRODUCTION

Cervical cancer is the fourth most common cancer among women worldwide in terms of both incidence and mortality rates^1^. Infection with high-risk human papillomavirus (HPV) contributes to almost all cervical cancer cases^2,3^. The oncogenic mechanism of HPV involves oncoproteins E6 and E7 which disrupt the normal cell lifecycle by deactivating tumour suppressor p53 and retinoblastoma protein (pRb), leading to uncontrolled cell proliferation, inappropriate cell survival and genomic instability^4,5^. Cervical squamous cell carcinoma (CSCC), the most common type, accounts for over 80% of these cases^6^. The introduction of prophylactic HPV vaccination and screening programs have significantly reduced the incidence rates of cervical cancer in developed countries ^2,7^. However, cervical cancer still disproportionately affects women in developing countries, where access to preventive measures and healthcare services is limited^6,8^. Staging remains a crucial factor influencing patient survival and is a key criterion for determining treatment^2,3^. Current treatment options for cervical cancer include surgery, chemotherapy, radiotherapy and combinations thereof, depending on the stage of the cancer, recurrence and other patient-specific factors^2^. In addition, immunotherapies have been approved and continue to be investigated for treating advanced, recurrent, or metastatic cervical cancers^9–11^.

Although HPV-positive CSCC is typically regarded as a single disease entity, the heterogeneous nature of cancer pathogenesis and of HPV infection result in significant disparities within this tumour type. The progression of HPV infection can vary, resulting in differences in mutation accumulation and epigenetic modifications. For instance, a study on The Cancer Genome Atlas (TCGA) cervical cancer (CESC) cohort reported an inconsistency in HPV integration event between HPV-16 and HPV-18^13^. The same study identified two subtypes (*i.e.*, keratin-high and keratin-low) of CSCC patients using an integrated analysis of multi-omics data. Gagliardi *et al.*^14^ reported unique patterns of genomic, epigenetic and pathway dysregulation associated with HPV clades. In particular, Chakravarthy *et al.*^12^ analysed the gene expression and DNA methylation profiles of CSCC patients from three international cohorts and identified two consensus molecular subtypes (CMSs) that correlate with patient survival. These studies have revealed heterogeneity in the molecular characteristics of HPV-positive CSCC patients, providing new insights into this disease and paving the way for the discovery of novel prognostic markers and therapeutic targets. However, the relationship between the CMS in CSCC tumours and their histology has not been extensively studied. Additionally, molecular analysis is typically expensive and slow, hindering its application in patient stratification and clinical trials.

Histological assessment of tissue sections is regarded as the gold standard for cancer diagnosis in clinical practice. H&E-stained tissue slides are widely available and contain rich information for clinical analysis. Recently, with the advancement of deep learning and computational pathology (CPath)^15–18^, several methods have been proposed to facilitate the analysis of multi-gigapixel whole slide images (WSIs) of H&E-stained tissue slides in tasks such as cell classification ^19,20^, nuclei detection and classification^21–23^, WSI classification^24–26^ and tumour-infiltrating lymphocytes (TILs) profiling^27,28^. In particular, many studies have explored the associations between molecular alterations and histological features presented in H&E-stained WSIs. For instance, researchers have demon-strated that various biomarkers determined by immunohistochemistry (IHC) or genomic analysis can be predicted from H&E WSIs using deep learning^29–32^. These studies indicate that the underlying genetic and epigenetic alterations in tumours can be reflected in histology, providing new tools and perspectives to investigate the relationships between genomic signatures and histological features in tissue sections.

In this study, we propose an end-to-end deep learning framework to explore the links between the histology patterns and two consensus molecular subtypes (*i.e.*, C1 and C2) of HPV-positive CSCC tumours identified by Chakravarthy *et al.*^12^. Our experiments on three international CSCC cohorts show that the CMS identified in Chakravarthy *et al.*^12^ can be predicted using routine H&E WSIs. Using two international CSCC cohorts, we demonstrate that our Digital-CMS scores are statistically significant in stratifying patients in relation to disease-specific survival (DSS) and disease-free survival (DFS), outperforming the CMS classification determined using DNA methylation data in Chakravarthy *et al.*^12^. Our extensive qualitative and quantitative histological analysis reveal distinct differences in the tumour microenvironment (TME) between C1 and C2 tumours, consistent with previous clinical findings. The contributions of this study are three-fold: First, compared to the molecular-determined CMS status, our Digital-CMS score derived from H&E WSIs is potentially more advantageous in clinical applications, saving the time and cost of molecular assays. Second, our TME analysis reveals statistically different immunological differences between C1 and C2 tumours, providing new insights into the different prognosis between C1 and C2 patients. Third, the exemplar histological patterns between C1 and C2 tumours can serve as surrogate visual markers by histopathology practitioners for patient prognostication or recruitment to clinical trials.

## RESULTS

### Pipeline for CMS prediction and TME profiling

We propose a deep learning framework to predict the genomically determined CMS from H&E-stained histology WSIs, as shown in Figure 1. Patches were first extracted from WSIs, and a domain-specific foundation model^17^ was used to extract their feature representations. The genomically determined CMS from Chakravarthy *et al.*^12^ was used to train our in-house TripletMIL model^33^, which was then used to generate both patch-level and a patient-level prediction of CMS status for an input WSI. The patient-level predictions were then also used as Digital-CMS scores for stratifying patients into different risk categories. Our patch-level predictions also identify representative tissue regions correlate with CMS status, providing valuable insights into the distinct TME patterns between CMSs. To identify visual patterns associated with CMS status, we perform clustering on representative WSI patches to select exemplar patches. To quantitatively profile TME differences between C1 and C2 tumours, two cell detection and classification models^21,34^ were first used to identify different types of cells in the representative tissue regions, and then cellular compositions and morphological features were extracted for a comprehensive TME profiling.

**Figure 1:**
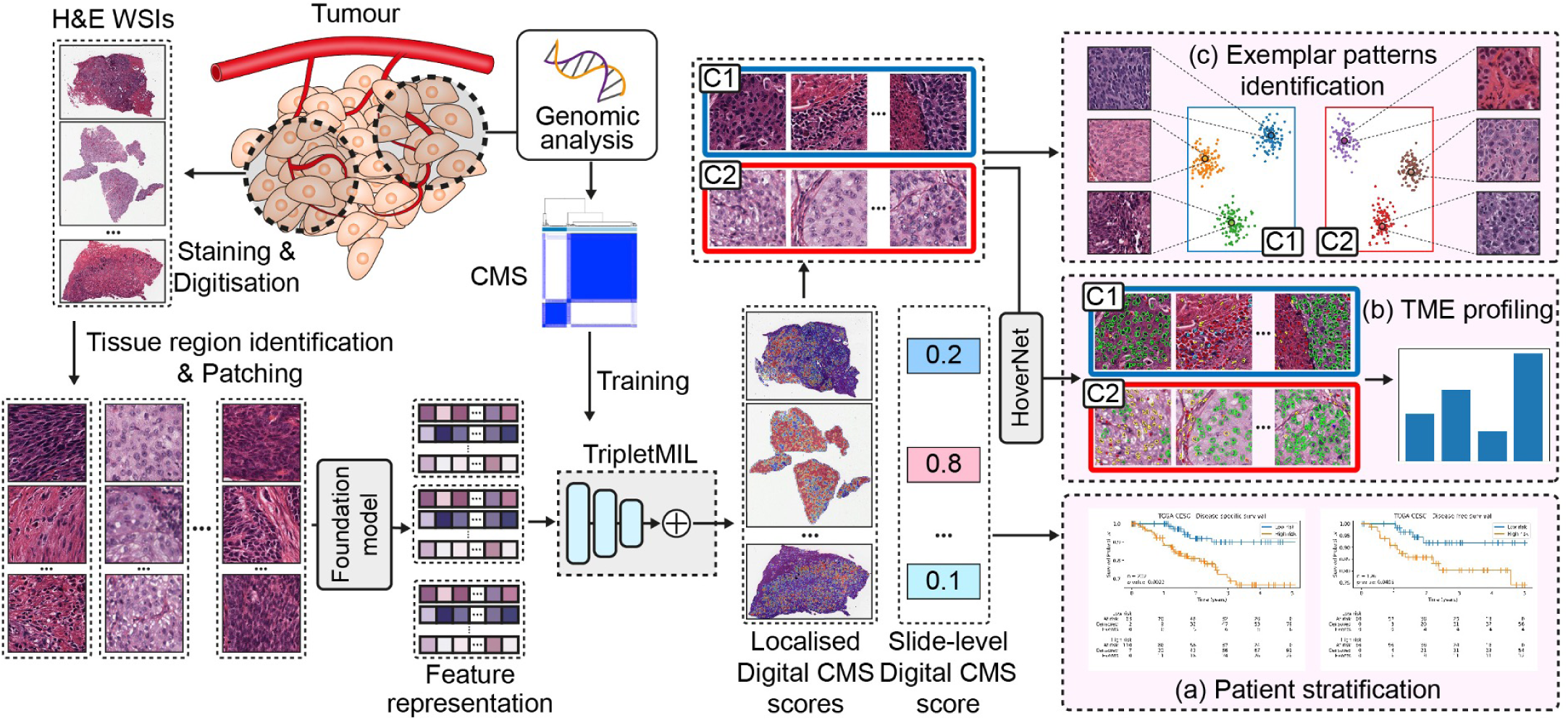
Proposed pipeline for CMS status prediction and TME profiling. Patches were extracted from the tissue regions of haematoxylin and eosin (H&E)-stained WSIs, the CMS was generated from the genomic analysis conducted by Chakravarthy *et al.* ^12^ and was used to train the TripletMIL. The trained TripletMIL model can generate localised Digital-CMS scores and a slide-level Digital-CMS score for an input WSI. (a) Slide-level Digital-CMS scores were used for patient stratification and survival analysis. (b) HoverNet was used to identify cells on the representative C1 and C2 patches, and statistical analysis was conducted to analyse the tumour microenvironment differences between C1 and C2 regions. (c) Clustering was performed to identify exemplar patterns from representative C1 and C2 patches.

### Prediction of CMS from H&E-stained WSIs

Our cross-cohort experiments show that the CMS (*i.e.*C1 and C2) generated from molecular analysis can be predicted from routine H&E histology images using the proposed algorithm. To evaluate the robustness and the generalisability of our proposed framework, we designed three cross-cohort settings for our analysis, as shown in Figure 2A. The three cohorts used in our study were collected from three different continents, reflecting demographic differences and data collection variations, such as staining and scanning variances. By conducting these cross-cohort validations, we attempt to ensure that our framework is not only effective across different populations but also resilient to the inherent variability of the digital histology images. In setting-1, our TripletMIL trained on Oslo-CSCC and Uganda-CSCC achieved an AUC of 0.78±0.03 on TCGA-CESC cohort; in setting-2, our TripletMIL trained on TCGA-CESC and Uganda-CSCC achieved an AUC of 0.85±0.04 on Oslo-CSCC cohort; in setting-3, our TripletMIL trained on TCGA-CESC and Oslo-CSCC achieved an AUC of 0.85±0.02 on Uganda-CSCC cohort, as shown in Figure 2B.

**Figure 2:**
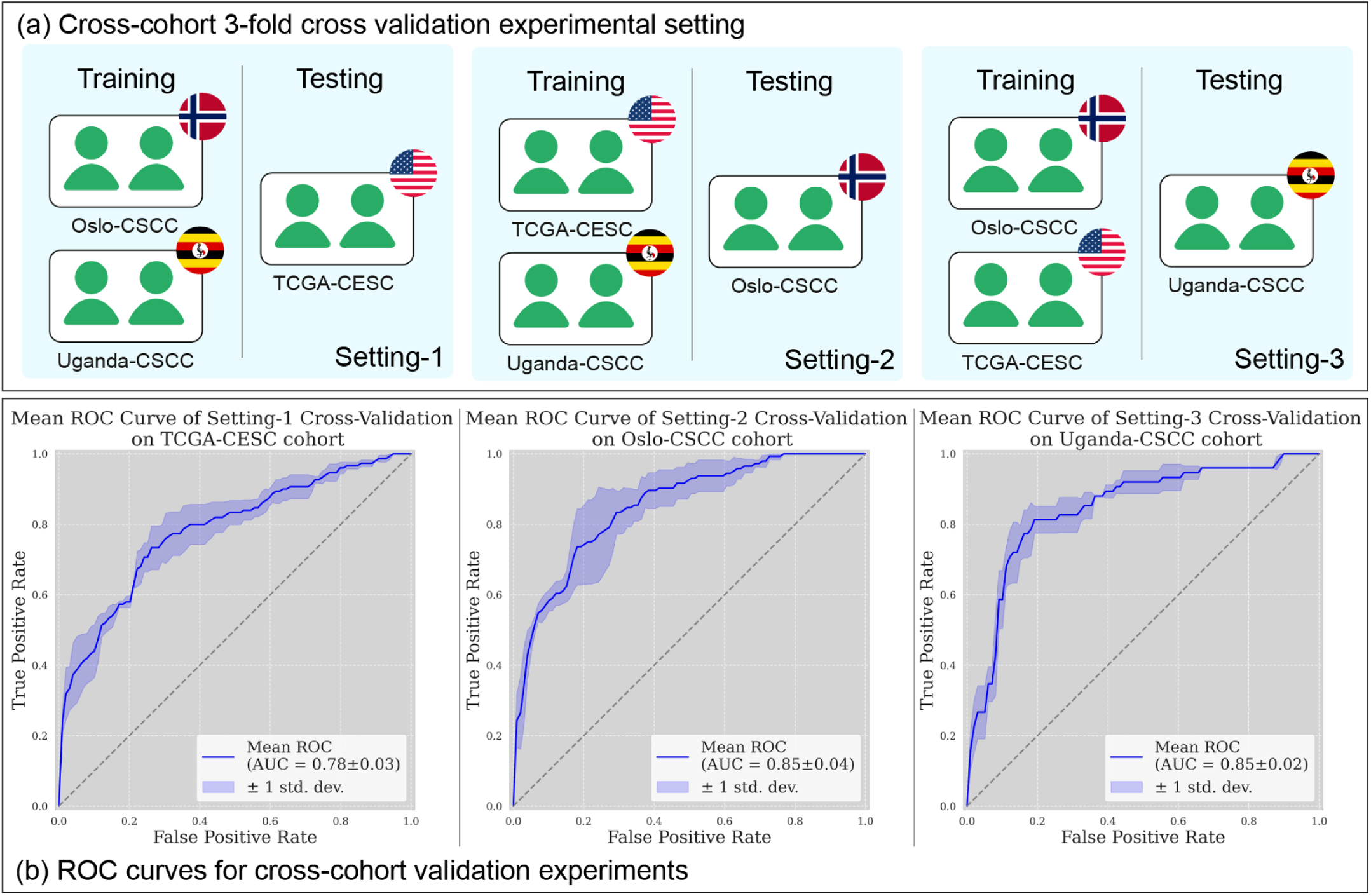
Cross-cohort experimental setting and results. (a) The three cross cohort experimental settings of our study. Each setting combines 2 cohorts for 3-fold cross validation experiments, and one held-out cohort for testing. (b) ROC curves of 3-fold cross validation experiments of our deep learning algorithms under the three settings. (CV: cross-validation)

### Digital CMS-score is associated with prognosis

Our experiments show that the proposed Digital-CMS scores can achieve statistically significant stratifications with DSS and DFS on TCGA-CESC and Oslo-CSCC cohort, under the cross-validation settings shown in Figure 2A. In setting-1, the proposed TripletMIL was trained on the combination of Oslo-CSCC and Uganda-CSCC cohorts, and Digital-CMS scores were then generated on the unseen TCGA-CESC cohort. Figure 3A shows significant prognostic differences between the low-risk group (predicted C1) and high-risk group (predicted C2) on TCGA-CESC cohort in terms of both DSS (p=0.0022) and DFS (p=0.0495). In setting-2, our TripletMIL was trained on the combination of TCGA-CESC and Uganda-CSCC cohorts, and Digital CMS scores were generated on the unseen Oslo-CSCC cohort. Figure 3C shows statistically significant prognostication between low-risk group (predicted C1) and high-risk group (predicted C2) on the Oslo-CSCC cohort in terms of both DSS (p=0.0495) and DFS (p=0.0282). In comparison, the CMS status determined using the DNA methylation data from Chakravarthy *et al.*^12^ does not produce statistically significant stratification for DSS (p=0.0975), as shown in Figure 3D. A high percentage of patients from the Uganda-CSCC cohort have human immunodeficiency virus (HIV) infection, with a 63% HIV+ rate. Given that HIV infection can significantly impact patient survival, this cohort is excluded from our survival analysis. As shown in Supplementary Table 1, HIV infection status is an independent predictor of overall survival on Uganda-CSCC cohort. The clinical characteristics, Digital-CMS score and Digital-CMS classifications of patients from TCGA-CESC and Oslo-CSCC cohort can be found in the Supplementary Data 2 and Supplementary Data 3, respectively.

**Figure 3:**
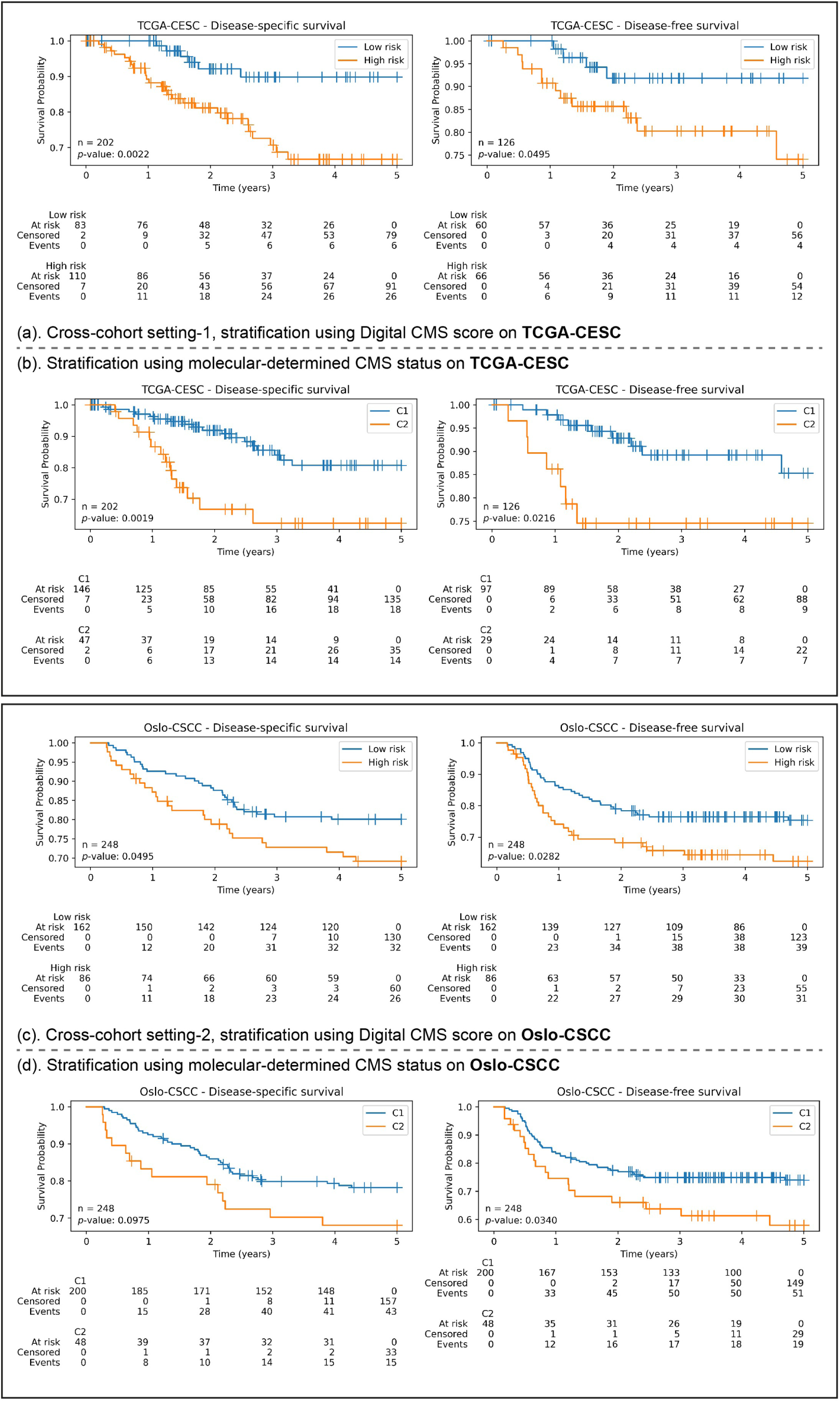
Survival analysis showing our digital-CMS score is able to stratify patients on DSS and DFS in the cross-cohort validations.

### Histological features associated with CMS across cohorts

Examining tissue regions associated with CMS status can provide valuable insights into histological features driven by molecular alterations and potentially serve as markers for determining patient prognosis. The patch-level CMS predictions from TripletMIL^33^ were used to identify localised morphological patterns unique to C1 and C2 tumours respectively. Figure 4A shows example WSI-level heatmaps highlighting the regions contributing to C1 and C2 predictions, respectively. One C1 and one C2 slide from each cohort were selected. Regions contributing more to C1 prediction are marked in blue, while regions contributing more to C2 prediction are marked in red. As can be seen from Figure 4A, a majority of the regions within C1 slides are marked in blue, indicating these regions are being identified as C1 regions, while a majority of C2 slides are marked in red, indicating these regions exhibit more C2-related features. To investigate what features are associated with C1 and C2 predictions, we identified exemplar histological patterns from representative C1 and C2 regions, as shown in Figure 4B. A consistent distinction of patterns between C1 and C2 can be observed across three cohorts. It can be observed that C1 regions generally have higher cellularity than the C2 regions; however, cells in C1 regions appear to be more regular in shape, while those in C2 often exhibit a higher degree of nuclear pleomorphism with enlarged nuclei.

**Figure 4:**
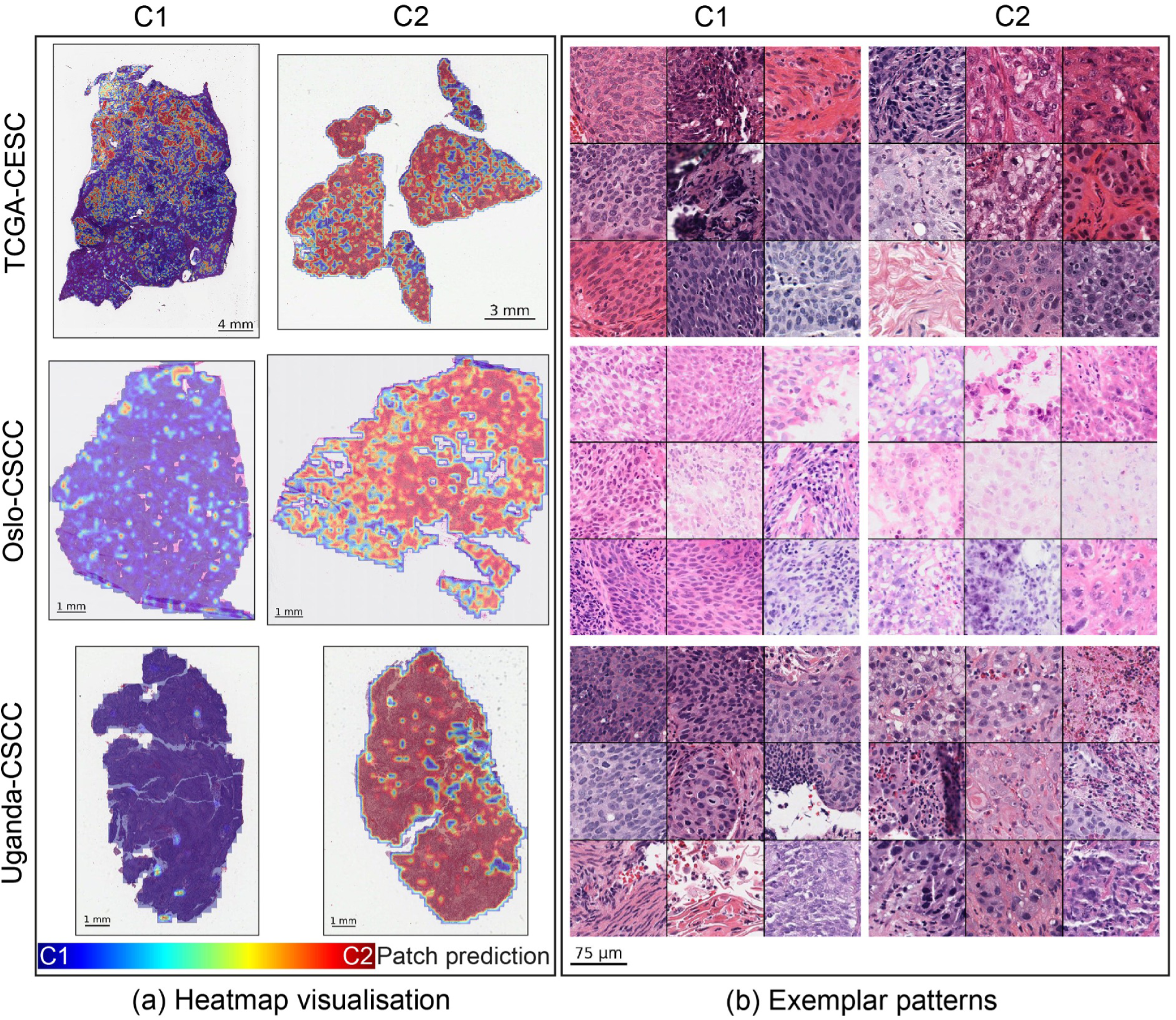
Visualisation of histological patterns associated with CMS across three cohorts. (a) WSI-level heatmap visualisation of one C1 slide and one C2 slide from each cohort; colour blue indicates higher prediction score towards C1, while colour red indicates higher prediction score towards C2. (b) Exemplar histological patterns identified from representative C1 and C2 regions in patients across three cohorts.

### TME patterns associated with CMS across cohorts

To quantitatively examine the TME patterns presented in the representative C1 and C2 regions, we first measured the cellular composition and morphological features of neoplastic, epithelial and connective cells. Our analysis showed that there exist distinct differences in these features between C1 and C2 tumours across three cohorts, as shown in Figure 5. It revealed a higher density of neoplastic and epithelial cells in C1 regions. The neoplastic and epithelial cells in C2 regions appear to be larger (measured by area) and more irregular (measured by the standard deviation of area) in size. This observation aligns with the visual examination of representative patches presented in Figure 4, indicating that C1 regions are generally densely populated with neoplastic and epithelial cell, while C2 regions contain more cells with enlarged nuclei, resulting in a lower cell density. In addition, we also found that the connective cells in C2 were more irregular in size (measured by the standard deviation of area and perimeter). We also examined these TME patterns in the entire sections and found that there were no significant difference across the three cohorts, indicating that these patterns are localised rather than uniformly distributed across the whole tissue section.

**Figure 5:**
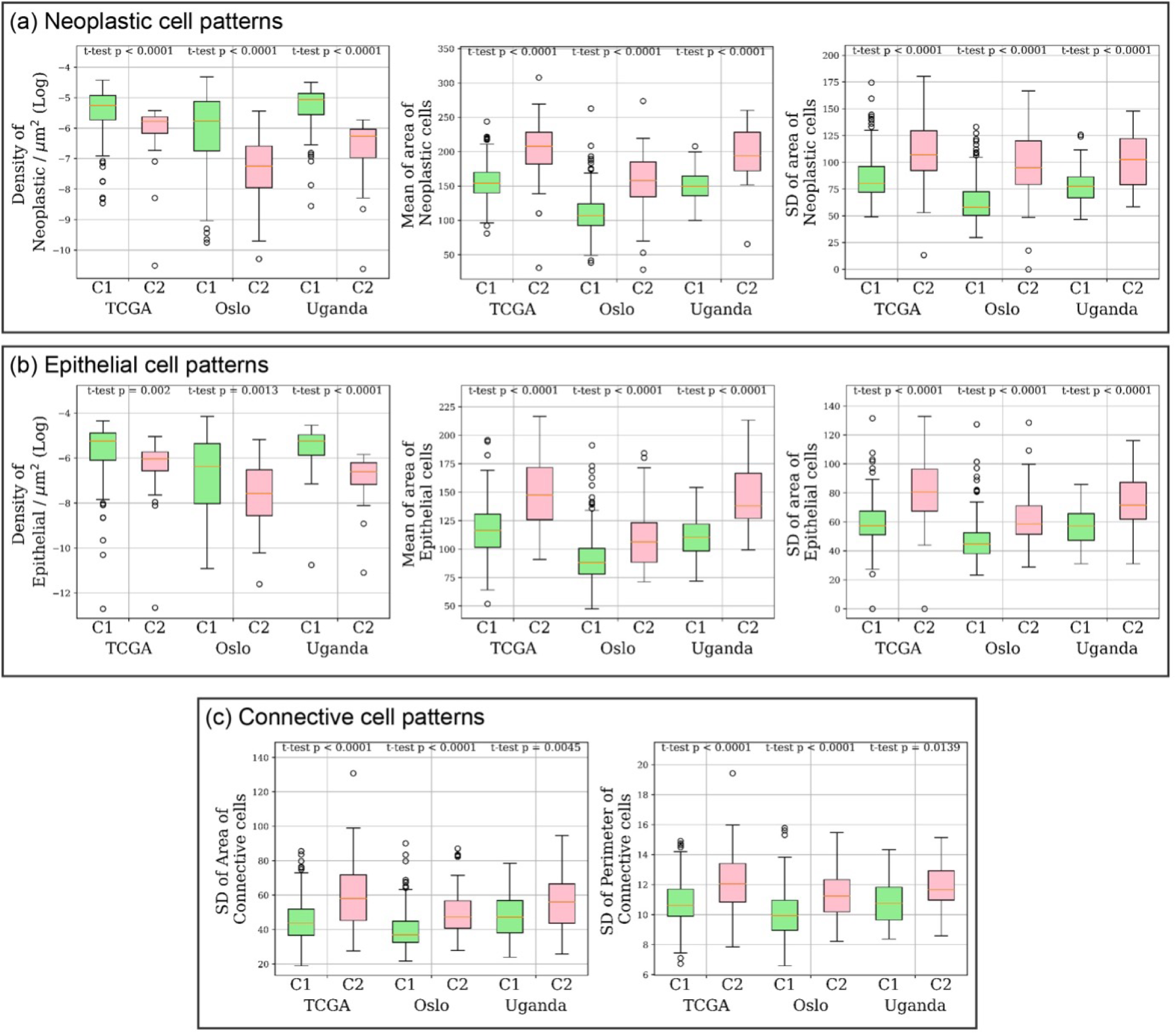
TME patterns associated with CMS across cohorts. Statistical analysis was conducted to compare the TME patterns between C1 and C2 regions identified by our algorithm. (a) TME patterns related to neoplastic cells, including the density, the mean and standard deviation (SD) of cells. (b) TME patterns related to epithelial cells, including the density, the mean and SD of area of cells. (c) TME patterns related to connective cells, including the SD of the area and perimeter of cells.

### Immunological patterns associated with CMS across cohorts

We analysed the density, ratio and the morphology of immune-related cells from representative C1 and C2 regions. As shown in Figure 6, C2 regions have an elevated level of neutrophil-to-lymphocyte ratio (NLR) and an elevated neutrophil density and ratio. These are in line with the cellular composition analysis conducted using DNA methylation data in Chakravarthy *et al.*^12^. We also noticed an elevated eosinophil ratio in C2 regions. In addition, C1 regions have a higher density of lymphocytes, and the size of lymphocytes (measured using the area and the perimeter of lymphocytes) appears to be larger in C1 regions.

**Figure 6:**
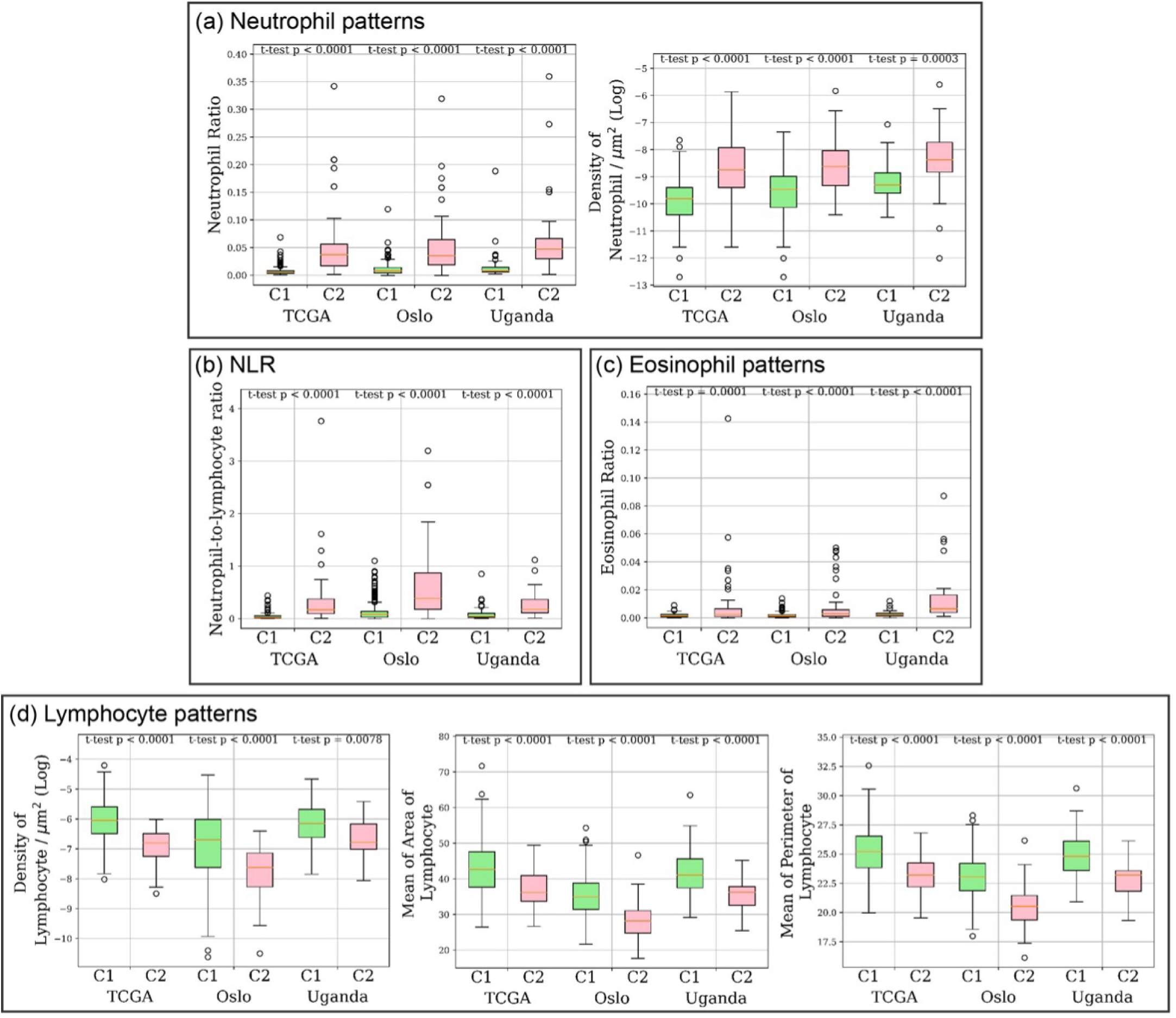
Immunological patterns associated with CMS across cohorts. Statistical analysis was conducted to compare the immunological patterns between C1 and C2 regions identified by our algorithm. (a) Immunological patterns related to neutrophil cells, including the ratio and density of cells. (b) Neutrophil-to-lymphocyte ratio (NLR) pattern. (c) Ratio of Eosinophil cells. (d) Immunological patterns related to lymphocyte cells, including the density, the mean of area and perimeter of cells.

We also examined the immunological patterns between C1 and C2 tumours on patient-level. A similar trend was observed, as shown in Figure 7. Elevated levels of the NLR and neutrophil ratio were noted in C2 patients, while C1 patients exhibited increased lymphocyte density, aligning with the findings of Chakravarthy *et al.*^12^. An elevated eosinophil ratio in C2 patients was observed and is consistent with the region-level analysis. In addition, we conducted a correlation analysis to examine the relationship between the IHC determined CD8+ score and our Digital-CMS score. A significant negative correlation was observed (p=0.0007), indicating that patients classified as C2 tend to have a lower CD8+ score.

**Figure 7:**
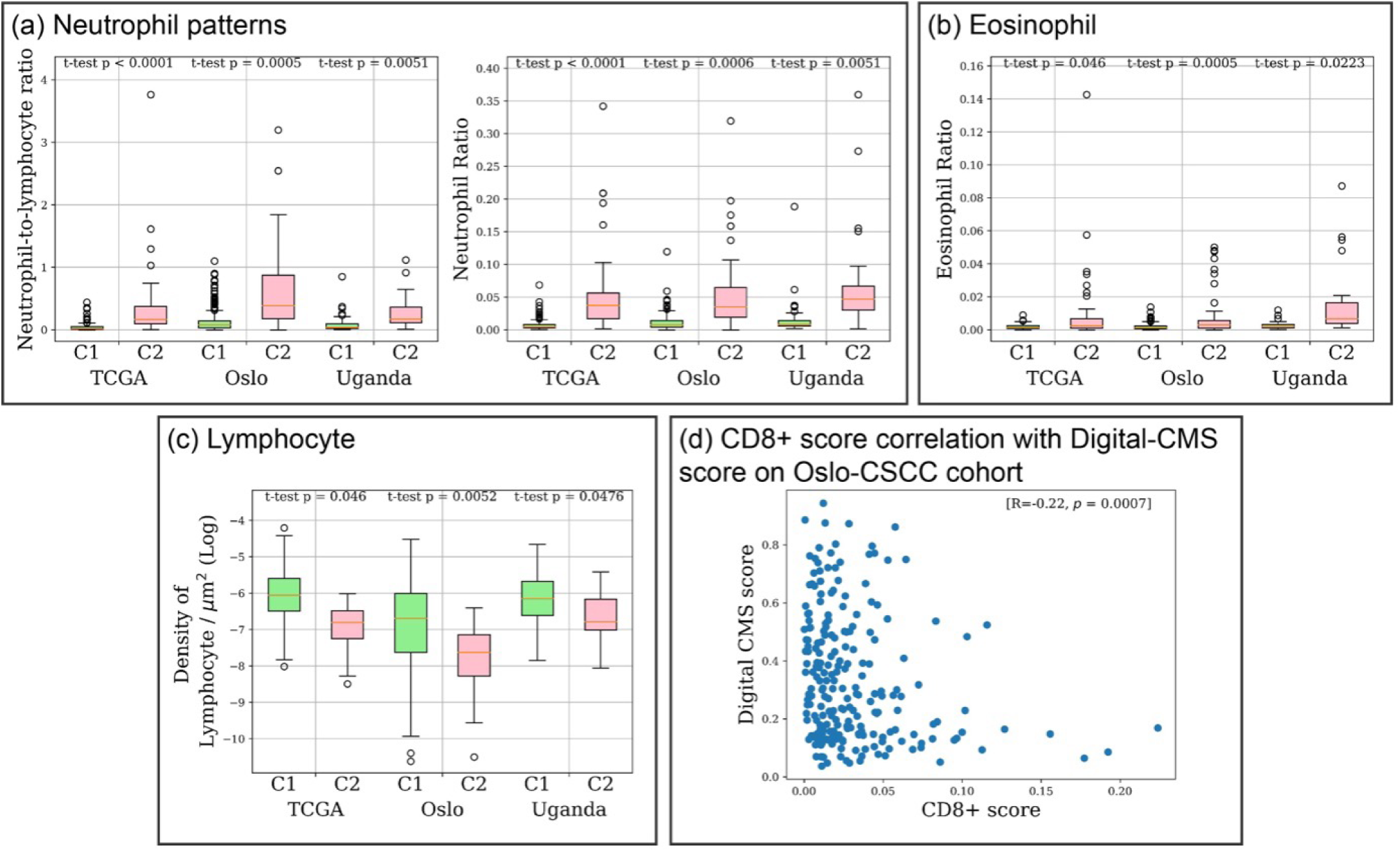
Immunological patterns associated with CMS across cohorts in patient-level. Statistical analysis was conducted to compare the immunological patterns between C1 and C2 patients. (a) Immunological patterns related to neutrophil cells, including the NLR and ratio of neutrophil. (b) Ratio of eosinophil. (c) Density of lymphocyte. (d) Correlation analysis between CD8+ score and Digital CMS score on Oslo-CSCC cohort.

### Higher lymphocytic infiltration in C1 tumours

TILs are an important biomarker in various types of cancers^28,35,36^ as they characterise the immune system’s interactions with tumours. Higher levels of TILs are associated with better clinical outcomes^35,37^ and can predict the efficacy of immunotherapies, such as immune checkpoint inhibitors^38^. We analysed the TILs pattern from H&E between C1 and C2 tumours by evaluating the lymphocyte density in tumour regions as well as its correlations with the Digital-CMS scores. As shown in Figure 8, across three cohorts, there is an elevated lymphocyte density in tumour regions in C1 tumours, as compared to C2 tumours. In addition, there is a negative correlation between Digital-CMS scores and lymphocyte density across three cohorts, indicating that as the likelihood of a patient being classified as C2 increases, the lymphocyte density decreases. This suggests that the lymphocytic infiltration is higher in C1 tumours, which may explain their better clinical outcomes.

**Figure 8:**
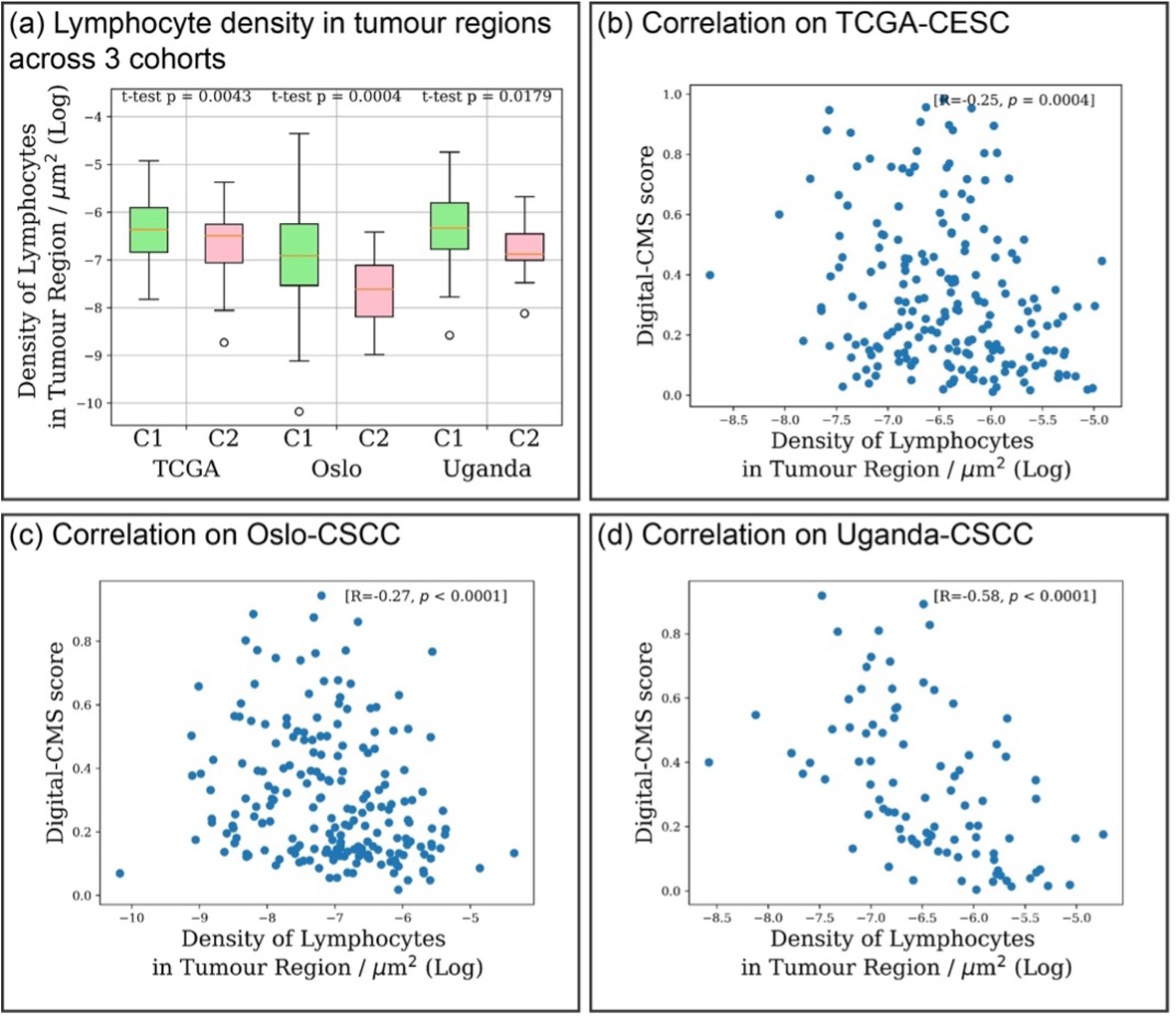
Different tumour infiltrated lymphocytes activity between C1 and C2 tumours. (a) C1 patients exhibit an elevated level of lymphocyte infiltration in tumour regions. (b) – (d) Negative correlations are shown between Digital-CMS scores and lymphocyte density in tumour regions across three cohorts.

## DISCUSSION

In this study, we proposed a deep learning framework to predict the consensus molecular sub-types of HPV-positive CSCC patients from digitised images of routine H&E-stained histology slides. Our model demonstrated strong predictive performance for CMS status across three international CSCC cohorts, validating its effectiveness and generalisability. This indicates that the molecularly determined CMS can be inferred from H&E-stained histology slides. In addition, our survival analyses show that our Digital-CMS scores were prognostically significant in terms of DSS and DFS on both TCGA-HNSC and Oslo-CSCC cohorts. We demonstrated that in the Oslo-CSCC cohort, Digital-CMS classifications using the proposed algorithm stratified patients more accurately in terms of DSS than the CMS classes obtained using DNA methylation data in Chakravarthy *et al.*^12^. The proposed algorithm offers a cost-effective and rapid alternative to genomic analysis for determining CMS status, providing a potential solution for clinical applications.

Our histological profiling of H&E-stained tissue slides revealed differences in cellular composition and morphology between C1 and C2 tumours. C1 representative regions were found to be more densely populated with regularly shaped tumour cells, with increased lymphocytic presence. C2 representative regions contained tumour cells with a higher level of nuclear pleomorphism, which may indicate higher grade of malignancy in C2 tumours. Our patient-level TME analysis showed that the CMS-associated TME patterns were localised rather than uniformly distributed throughout the tissue. This highlights the importance of regional analysis with the help of our algorithm in understanding the tumour microenvironment. However, establishing the linkage between nuclear shape irregularity and specific genetic and epigenetic alterations is challenging^39,40^. Further investigations are needed to examine the relationships between histological features and CMS. It is worth noting that despite significant staining differences between the three cohorts, as shown in Figure 4, the foundation model^17^ used in this study and our TripletMIL were able to discern consistent histological correlates with the CMS status.

Our immunological profiling identified an elevated NLR, neutrophil ratio and density in C2 tumours, which are consistent with the findings of Chakravarthy *et al.*^12^ which was performed using DNA methylation data. In addition, our analysis based on H&E histology identified an enlarged lymphocyte cell size in C1 tumours, which may indicate the presence of activated lymphocytes, as an increase in size of activated lymphocytes has been reported in previous literature ^41,42^. Higher density of lymphocytes in tumour-rich areas in C1 tumours indicates a higher lymphocytic infiltration in C1 patients. In addition, correlation analysis on Oslo-CSCC cohort between CD8+ score and Digital-CMS score also showed a higher presence of CD8+ T cells in C1 patients, which may explain their better prognosis. Correlation of tumours with an elevated eosinophil presence with worse immune-response and a poor prognosis in those tumours has also been reported in previous studies^43,44^, which may explain our finding of a higher eosinophil ratio in C2 tumours.

With the advancement of immunotherapy, several studies^10,45,46^ have demonstrated that immune checkpoint inhibitors can enhance the immune response and improve patient outcomes in cervical cancer patients. In particular, the combination of pembrolizumab (Keytruda, Merck) with chemoradiotherapy has been approved by the US FDA for treating advanced stage cervical cancer patients^47^. However, analyses of the tumour microenvironment in both Chakravarthy *et al.*^12^ and this study indicate significant differences between the immune microenvironments of C1 and C2 tumours. The relatively low level of immune infiltration in C2 tumours suggests they may respond poorly to immunotherapy. Therefore, further stratification of cervical cancer patients is necessary to investigate their response to immunotherapy in future clinical trials, with the Digital-CMS score proposed in this work serving as a potential biomarker for this stratification.

Our study has some limitations. First of all, our algorithm’s prediction accuracy on TCGA-CESC is not as good as that on the Oslo-CSCC or Uganda-CSCC cohorts. However, the Digital-CMS score is still able to achieve statistically significant stratification on TCGA-CESC. This may indicate that there exist discrepancies between the CMS of TCGA-CESC and other cohorts, since the method used to determine CMS on these cohort is different in Chakravarthy *et al.*^12^. Therefore, validations on larger cohorts may be needed to further validate these subtypes and their histological correlations. In addition, the TCGA-CESC and Oslo-CSCC cohorts used in our study were curated from high-income countries where HPV prevention measures are widely accessible. Consequently, these cohorts may not accurately represent patients from middle-/low-income countries where access to prevention or medical care is limited. Therefore, large-scale validation studies on cohorts from these regions are necessary to further validate the consensus subtypes and ensure their applicability across diverse populations.

Finally, our study has revealed histological associations of the genomically-derived CMS proposed by Chakravarthy *et al.*^12^. This further supports the existence of molecular and histological heterogeneities within HPV-positive CSCC tumours. Despite advancements in prevention and screening methods, cervical cancer continues to pose a significant health challenge, especially in developing countries^2^. This underscores the critical need for identifying more effective prognostic biomarkers and therapeutic targets to improve patient outcomes. By enhancing our understanding of the tumour microenvironment and its variations, our research contributes to the ongoing efforts to develop tailored treatment strategies and improve prognosis for patients with HPV-positive CSCC.

## METHODS

### Ethics statement

The ethical approval for Oslo-CSCC cohort was obtained from the Regional Committee for Medical Research Ethics South East Norway (2016/2179). Written informed consent was achieved from all patients in Oslo-CSCC cohort. No further ethical approval was required for TCGA-CESC and Uganda-CSCC cohorts because the tissue slides for both cohorts are publicly available for research purposes.

### Data collection

TCGA-CESC cohort consists of 203 cases collected from 27 medical centres in the USA. Uganda-CSCC cohort consists of 94 cases collected from the Uganda Cancer Institute. The formalin-fixed, paraffin-embedded (FFPE) H&E-stained WSIs of these two cohorts were retrieved from the GDC Data Portal (https://portal.gdc.cancer.gov/) under the project TCGA-CESC and CGCI-HTMCP-CC, respectively.

Oslo-CSCC cohort consists of 248 cases collected from the Oslo University Hospital. H&E-stained histology slides of Oslo-CSCC cohort were scanned using an Olympus VS200 scanner (Olympus Corporation) at a resolution of 0.2725 microns per pixel (mpp). The bfconvert tool from Bio-Formats library^48^ was used to convert WSIs of Oslo-CSCC cohort from Olympus VSI format (.vsi) format to Generic TIFF (.tiff) format for further processing.

The clinical data of TCGA-CESC was retrieved from the study by Liu *et al.*^49^. The clinical data of Oslo-CSCC was collected from Oslo University Hospital. It is worth noting that the clinical follow-up data for the Oslo-CSCC cohort used in this study differs slightly from that used in Chakravarthy *et al.*^12^ due to updated patient follow-up information.

The TCGA-CESC cohort includes patients with an age range of 21-85 years and a median age of 47. The Oslo-CSCC cohort includes patients with an age range of 22-82 years and a median age of 54. The Uganda-CSCC cohort includes patients with an age range of 26-82 years and a median age of 45.

### Consensus molecular subtype determination

The CMS status used in this study was retrieved from the study by Chakravarthy *et al.*^12^. Based on the original study, the CMS status of TCGA-CESC cohort was determined from consensus clustering on the top 10% most variably expressed genes; the CMS status of Oslo-CSCC and Uganda-CSCC cohorts was inferred from the DNA methylation data with a support vector machine (SVM) trained on the TCGA-CESC cohort.

### Pre-processing of WSIs

Tissue mask of each WSI was generated using TIAToolbox^50^ by identifying tissue regions and removing background and artifacts. Then, sliding window patching implemented by TIAToolbox^50^ was used to generate non-overlapping image patches of size 256×256 at a spatial resolution of 0.5 mpp from tissue regions. Image patches were saved if the tissue portion is greater than 80%, based on the generated tissue mask.

### Feature representations of image patches

Foundation model UNI^17^ was used to generate feature representations of image patches from the previous step. UNI is a domain-specific foundation model for H&E-stained histology images. It is developed based on ViT structure^51^ and was trained on an in-house dataset Mass-100K^17^ using DINOv2^52^. The Mass-100K dataset comprises 100,130,900 images from 100,426 histology slides across 20 tissue types, collected from Massachusetts General Hospital (MGH), Brigham and Women’s Hospital (BWH), and the Genotype-Tissue Expression (GTEx) consortium. The three cohorts used in this study were not seen by UNI since it was not trained on any publicly available datasets such as TCGA. Therefore, the feature representations from UNI would not have bias towards any dataset used.

### Bag construction for multiple instance learning

Since only the patient-level CMS status is available, we adopt the multiple-instance learning (MIL) paradigm for CMS status prediction. In a MIL setting, each patient is treated as a bag of patches, and the patient-level CMS status is given to each bag as the ground-truth label. Let *B* represent a set of patches extracted from a single slide, we consider *{B*_1_*, B*_2_*, . . ., B_N_ }* as *N* bags of patches extracted from *N* slides. The corresponding ground-truth labels *Y_n_* ∈ *{*0, 1*}* for *n* = 1*, . . ., N* denote the slide-level CMS label, where 0 represents C1 and 1 represents C2. Each bag *B* contains different number of patches *{I*_1_*, I*_2_*, . . ., I_K_}, K* ∈ ℤ^+^, where each patch is represented by a 1024-dimensional feature vector *f ∈* ℝ^1024^ generated by UNI.

### C1/C2 prediction using TripletMIL

TripletMIL^33^ is a ranking-based MIL framework for biomarker prediction. TripletMIL takes the feature representations of patches from a patient, and outputs the Digital-CMS scores at both patch-level and patient-level. In the TripletMIL training process, a bag of feature representations *{f*_1_*, f*_2_*, . . ., f_K_}*, *K ∈* ℤ^+^*, f ∈* ℝ^1024^ is input into a multi-layer perceptron (MLP), which learns the feature representations and generates a Digital-CMS score for each patch. An average aggregation is used to generate a patient-level Digital-CMS score for each patient. TripletMIL uses a triplet loss function to learn the correct ranking between samples. Triplet loss aims to establish the correct order among samples and to learn precise similarity relationships between them. In each iteration of the triplet ranking training, two different C1 samples and one C2 sample were randomly drawn from the training set, and a trainable MLP and an average aggregation are used to generate a patient-level Digital-CMS score for each sample. The three scores are used to calculate the triplet loss *L*_tri_, which is formulated as follows:

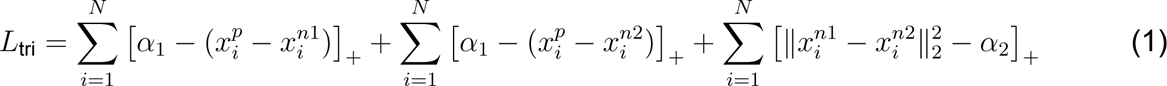

where 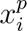 denotes the score for the C2 sample, and 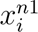 and 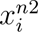 denote the scores of the two different C1 samples. *α*_1_ and *α*_2_ are the margin parameters for constraining inter-class variation and intra-class variation respectively. In our experiments, *α*_1_ is set to 0.5 and *α*_2_ is set to 0.1. *L*_tri_ imposes two types of constraints over three input samples. The first two terms consider the differences of the predicted values between the C2 and two C1 samples. It forces the model to predict a higher value for C2 samples than any of the C1 samples by at least *α*_1_. The third term aims to reduce the distances among C1 samples to reduce the intra-class variation.

### Model training and validation

We performed 3-fold stratified cross validation (CV) on the three cohorts used in our study in three settings, as shown in Figure 2A. In each setting, two cohorts were combined to form a combined set for 3-fold CV, and the left-out cohort is used as the testing set. On the combined set, data from two cohorts were mixed and split into 3 folds stratified based on CMS status; in each experiment, 2 folds of data were selected for training the model, and the left-out fold was used for training monitoring and saving the best-performing model. We trained the model for 20 epochs with early stopping when the training does not produce a positive loss. Learning rate was set to 3 *×* 10*^−^*^3^. Stochastic gradient descent (SGD) was used as optimiser, momentum was set to 0.9 and weight decay was set to 1*×*10*^−^*^4^. Learning rate was decayed by 0.1 every 10 epochs. The training code was implemented in PyTorch. One NVIDIA Tesla V100 on a NVIDIA DGX 2 was used for training the model.

### Survival analysis with Digital-CMS score

Survival analysis was conducted on Setting-1 and Setting-2 to test the prognostic value of our Digital-CMS scores. For each setting, the best performing model from 3-fold CV was selected based on the performance on the validation fold. Then, the Digital-CMS score for each patient on the left-out cohort was generated. For survival analysis, in each setting, the combined cohort was used as the discovery set, and the left-out cohort was used as the validation set. Cut-off value for stratifying patients into high-risk (predicted C2) and low-risk (predicted C1) groups were selected based on the ROC curve of the best performing model on the discovery set. Specifically, the cut-off value was determined using Youden’s J statistic^53^ with the goal of maximising the sum of sensitivity and specificity at a chosen point on the ROC curve. The Kaplan-Meier curves for DSS and DFS were plotted, and the significance in survival differences between high and low risk groups were tested using the log-rank test.

### Histological feature identification

Histological feature identification was performed to identify the unique patterns reflected associated with CMS status from H&E-stained tissue slides. Patients were divided to C1 and C2 groups based on their CMS status, and the best performing model was used to generate a patch-level Digital-CMS score for each image patch. For C1 patients, we select 20 patches with the highest predicted probability of being C1 (lower Digital-CMS scores); for C2 patients, we select 20 patches with the highest predicted probability of being C2 (higher Digital-CMS scores). Within each group, k-means clustering was used to cluster the selected patches into 9 clusters using the deep feature representations generated by UNI^17^, and 9 histological patterns of each group were identified.

### Cellular composition and statistical analysis

Cellular composition analysis was performed to identify the unique tumour-microenvironment and immunological patterns between C1 and C2 tumours. HoverNet^21^ trained on PanNuke dataset^23^ and AugHoverNet^34^ trained on CoNIC dataset^22^ was used to detect and classify nuclei from H&E-stained WSIs. The nuclei detection and segmentation was performed at WSI level with TIAToolbox’s pipeline^50^ to prevent classification inaccuracies associated with nuclei located at the edges of image patches. HoverNet-PanNuke classifies nuclei into 5 categories: neoplastic, inflammatory, connective, necrosis and non-neoplastic; while AugHoverNet-CoNIC classifies nuclei into 6 categories: neutrophil, epithelial, lymphocyte, plasma, eosinophil and connective. For localised analysis, representative C1 and C2 patches were selected using the method described in the above section, and different features were calculated based on the nuclei detection results on these patches. Specifically, for each cell type, cell density was calculated as the number of cells (*N*) divided by the area (*A*) of the patches (*A* in *µm*^2^), using the formula: 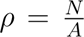. The natural logarithm (*ln*) was applied to the cell density values to facilitate data analysis and interpretation. The cell ratio is calculated as the number of cells in question divided by the total number of cells detected in the region. The neutrophil-to-lymphocyte ratio is calculated as the number of neutrophils detected divided by the number of lymphocytes detected in the region. The morphological features of each nucleus, including area and perimeter, were calculated using OpenCV based on the contours of the nuclei identified by HoverNet and AugHoverNet. Mean, median and standard deviation are used to measure different statistics of each morphological feature of a cell type from a representative region. The lymphocytic density in tumour regions is calculated using the formula stated above, the tumour region is defined as any patch with a size of 128 *×* 128 *µm*^2^ containing more than 50 neoplastic cells. For WSI-level analysis, the features were calculated using the aforementioned method applied to the entire WSI. To compare the differences between C1 and C2 groups, box plots were used to visualise the distribution of the data, and independent samples two-tailed t-test was conducted to statistically assess the significance of the differences. Given the large sample size, normality was assumed based on the central limit theorem. However, the Mann-Whitney U test and Cliff’s Delta were also calculated to assist in interpreting the results. Detailed results of the statistical tests can be found in the Supplementary Data 4-15. The Spearman correlation test was used to evaluate the relationship between Digital-CMS scores and biological features (*i.e.*, the lymphocyte density in tumour regions, the CD8+ scores) across three cohorts.

### CD8+ scoring

Immunohistochemistry was performed using monoclonal mouse anti-CD8 (1:150, clone 4B11, NCL-CD8-4B11, Novocastra, Leica Microsystems, Newcastle Upon Tyne, UK) on sections from paraffin embedded tumour samples. Antigen retrieval was performed with PT-link and Envision target retrieval solution (Dako, Glostrup, Denmark), stained by the DAKO Envision Flex+ system (Dako), and visualised in brown by 3,3-diaminobenzidine (DAB) and blue (haematoxylin) as described Chakravarthy *et al.*^12^. Digital Images were acquired by a NanoZoomer-XR slide scanner (Hamamatsu, Hamamatsu City, Japan). The sections were evaluated for CD8-positive cells using DAB staining. The sections were digitised at 20*×* magnification and were subsequently analysed at a reduced scale of 50%, providing a resolution of 0.92 mpp. Colour deconvolution was used to extract the staining intensities of the DAB colour channel. Threshold segmentation was employed to generate binary maps distinguishing areas positive for CD8 expression. To quantify the prevalence of CD8-positive cells, an area fraction was calculated as the ratio of the CD8-positive stained areas relative to the overall tissue biopsy area.

## ACKNOWLEDGEMENTS

R.W. is grateful to Dr. Muhammad Dawood (University of Oxford, UK) and Dr. Quoc Dang Vu (Histofy Ltd., UK), both formerly at the TIA Centre of the University of Warwick, for discussions and feedback on the topic of this study. This study was partly supported by a PhD studentship to R.W. funded by the General Charities of the City of Coventry (grant reference: 67148). R.W. is also grateful to the Computer Science Doctoral Training Centre at the University of Warwick for providing matched funding for this research. The results published here are in part based on data generated by the The Cancer Genome Atlas (TCGA) Research Network: https://www.cancer.gov/tcga.

## AUTHOR CONTRIBUTIONS

**R.W.**: Conceptualization, Methodology, Software, Validation, Formal analysis, Investigation, Data Curation, Writing - Original Draft, Writing - Review & Editing, Visualization, Project administration. **G.N.G.**: Software, Formal analysis, Investigation, Resources, Data Curation, Writing - Review & Editing, Visualization. **V.E.S.**: Methodology, Validation, Formal analysis, Investigation, Resources, Data Curation, Writing - Review & Editing. **K.A.F.V.**: Validation, Investigation, Resources, Data Curation, Writing - Review & Editing. **H.L.**: Methodology, Validation, Formal analysis, Investigation, Resources, Data Curation, Writing - Review & Editing, Supervision. **L.S.Y.**: Conceptualization, Methodology, Validation, Formal analysis, Writing - Review & Editing, Supervision, Funding acquisition. **N.R.**: Conceptualization, Methodology, Validation, Formal analysis, Writing - Review & Editing, Supervision, Funding acquisition

## COMPETING INTERESTS

N.R. is the Chief Executive Officer, Chief Scientific Officer and a Director of Histofy Ltd. The authors declare no other competing interests.

## DATA & CODE AVAILABILITY

TCGA-CESC and Uganda-CSCC cohorts can be accessed at GDC Data Portal (https://portal.gdc.cancer.gov/). Oslo-CSCC cohort can be accessed upon reasonable request, and with additional ethical approvals. Source code for this paper can be found at https://github.com/ruoyussh/cervical-cms.

