## Supplementary material for "A Deep Learning Framework for Predicting Prognostically Relevant Consensus Molecular Subtypes in HPV-Positive Cervical Squamous Cell Carcinoma from Routine Histology Images": Supplementary Information.pdf

**Supplementary Data Titles:**

**Supplementary Data 1:** Multivariate analysis of risk factors on Uganda-CSCC cohort on overall survival.

**Supplementary Data 2:** Clinical characteristics of TCGA-CESC cohort. Digital-CMS classification was derived from Digital-CMS score using cutoff value of 0.2167, selected on the discovery cohort using Youden's J statistic.

**Supplementary Data 3:** Clinical characteristics of Oslo-CSCC cohort. Digital-CMS classification was derived from Digital-CMS score using cutoff value of 0.381, selected on the discovery cohort using Youden's J statistic.

**Supplementary Data 4:** Statistical results of differences between C1 and C2 representative regions on cellular features identified with HoverNet-Pannuke, on TCGA-CESC cohort.

**Supplementary Data 5:** Statistical results of differences between C1 and C2 representative regions on cellular features identified with HoverNet-Pannuke, on Oslo-CSCC cohort.

**Supplementary Data 6:** Statistical results of differences between C1 and C2 representative regions on cellular features identified with HoverNet-Pannuke, on Uganda-CSCC cohort.

**Supplementary Data 7:** Statistical results of differences between C1 and C2 representative regions on cellular features identified with AugHoverNet-Conic, on TCGA-CESC cohort.

**Supplementary Data 8:** Statistical results of differences between C1 and C2 representative regions on cellular features identified with AugHoverNet-Conic, on Oslo-CSCC cohort.

**Supplementary Data 9:** Statistical results of differences between C1 and C2 representative regions on cellular features identified with AugHoverNet-Conic, on Uganda-CSCC cohort.

**Supplementary Data 10:** Statistical results of differences between C1 and C2 tumours (WSI-level) on cellular features identified with HoverNet-Pannuke, on TCGA-CESC cohort.

**Supplementary Data 11:** Statistical results of differences between C1 and C2 tumours (WSI-level) on cellular features identified with HoverNet-Pannuke, on Oslo-CSCC cohort.

**Supplementary Data 12:** Statistical results of differences between C1 and C2 tumours (WSI-level) on cellular features identified with HoverNet-Pannuke, on Uganda-CSCC cohort.

**Supplementary Data 13:** Statistical results of differences between C1 and C2 tumours (WSI-level) on cellular features identified with AugHoverNet-Conic, on TCGA-CESC cohort.

**Supplementary Data 14:** Statistical results of differences between C1 and C2 tumours (WSI-level) on cellular features identified with AugHoverNet-Conic, on Oslo-CSCC cohort.

**Supplementary Data 15:** Statistical results of differences between C1 and C2 tumours (WSI-level) on cellular features identified with AugHoverNet-Conic, on Uganda-CSCC cohort.
